# *Caenorhabditis elegans* germ granules are present in distinct configurations that differentially associate with RNAi-targeted RNAs

**DOI:** 10.1101/2023.05.25.542330

**Authors:** Celja J. Uebel, Sanjana Rajeev, Carolyn M. Phillips

## Abstract

RNA silencing pathways are complex, highly conserved, and perform widespread, critical regulatory roles. In *C. elegans* germlines, RNA surveillance occurs through a series of perinuclear germ granule compartments—P granules, Z granules, SIMR foci, and *Mutator* foci—multiple of which form via phase separation and exhibit liquid-like properties. The functions of individual proteins within germ granules are well-studied, but the spatial organization, physical interaction, and coordination of biomolecule exchange between compartments within germ granule “nuage” is less understood. Here we find that key proteins are sufficient for compartment separation, and that the boundary between compartments can be reestablished after perturbation. Using super-resolution microscopy, we discover a toroidal P granule morphology which encircles the other germ granule compartments in a consistent exterior-to-interior spatial organization. Combined with findings that nuclear pores primarily interact with P granules, this nuage compa rtment organization has broad implications for the trajectory of an RNA as it exits the nucleus and enters small RNA pathway compartments. Furthermore, we quantify the stoichiometric relationships between germ granule compartments and RNA to reveal discrete populations of nuage that differentially associate with RNAi-targeted transcripts, possibly suggesting functional differences between nuage configurations. Together, our work creates a more spatially and compositionally accurate model of *C. elegans* nuage which informs the conceptualization of RNA silencing through different germ granule compartments.

## INTRODUCTION

Thousands of transcripts undergo RNA surveillance to ensure proper gene expression in *C. elegans*. Transcript monitoring and silencing is performed by conserved regulatory small RNAs comprised of three distinct branches: microRNAs (miRNA), piwi-interacting RNAs (piRNA), and small interfering RNAs (siRNA). Each small RNA branch differs in biogenesis, protein interactors, transcript targets, and primary mechanisms of action, but the fundamental means of RNA surveillance are conserved: small RNAs, ranging in size from 18-30 nucleotides, are bound by Argonaute proteins and together target fully or partially complementary RNAs to direct RNA silencing through transcriptional or post-transcriptional mechanisms (reviewed in (Ketting et al., 2021)).

Small RNA-directed silencing is efficient. Exogenously introduced double-stranded RNAs are routed through the siRNA pathway and can silence transcripts to biochemically undetectable levels (Fire et al., 1998). A reduction in transcript levels can be detected by 4 hours and silencing outcomes can be phenotypically observed by 24 hours (Kamath et al., 2000; Ouyang and Seydoux, 2022). Silencing signals directed at germline-expressed genes can also be inherited through multiple generations, mediated by siRNA amplification, maternal and paternal small RNA inheritance, and small RNA-directed accumulation of repressive chromatin marks on targeted loci (Alcazar et al., 2008; Ashe et al., 2012; Burkhart et al., 2011; Grishok et al., 2000; Shirayama et al., 2012). RNA silencing also functions to ensure proper gene expression and protect against deleterious transcripts. In brief, 1) miRNAs regulate genes involved in cell patterning, development, and lifespan (reviewed in (Ambros and Ruvkun, 2018)), 2) piRNAs engage most germline genes and can target transposons, including the Tc3 transposon (Bagijn et al., 2012; Batista et al., 2008; Das et al., 2008; Lee et al., 2012; Shen et al., 2018), and 3) endogenous siRNAs target additional transposons, gene duplications, pseudogenes, germline genes, and repetitive elements (Fischer et al., 2011; Gu et al., 2009). “Primary” small RNAs, including piRNAs and a subset of siRNAs, feed into a downstream amplification step which produces “secondary” siRNAs. The amplification of primary small RNAs to secondary siRNAs creates a robust, heritable, efficient silencing signal (Aoki et al., 2007; Grishok et al., 2000; Pak and Fire, 2007; Vasale et al., 2010). Together, small RNA pathways accomplish widespread regulatory roles and are required for proper development and fertility.

In the germline, the subcellular organization of small RNA pathways occurs within perinuclear germ granules. At least four distinct compartments, collectively referred to here as *nuage*, facilitate efficient RNA surveillance, coordinate small RNA amplification, and ensure small RNA inheritance (Manage et al., 2020; Phillips et al., 2012; Pitt et al., 2000; Sheth et al., 2010; Wan et al., 2018). P granules, the first discovered *C. elegans* nuage compartment, contain both nascent transcripts and key small RNA pathways components including the Argonautes PRG-1, ALG-3, CSR-1, and WAGO-1 (Batista et al., 2008; Claycomb et al., 2009; Conine et al., 2010; Gu et al., 2009; Sheth et al., 2010). P granules also contain other small RNA associated factors including the <u>D</u>icer-related <u>h</u>elicase DRH-3 and the <u>R</u>NA-<u>d</u>ependent <u>R</u>NA <u>p</u>olymerase (RdRP) EGO-1, which act together to produce a subset of secondary siRNAs (Claycomb et al., 2009; Gu et al., 2009). Also localized to P granules are the <u>p</u>iRNA-induced silencing-<u>d</u>efective, PID-1, and the RNase, PARN-1, which are required for the synthesis and processing of piRNAs (Cordeiro Rodrigues et al., 2019; de Albuquerque et al., 2014; Tang et al., 2016). Further, most nuclear pores (75%) associate with P granules, indicating that the majority of nascent transcripts enter P granules upon nuclear export (Pitt et al., 2000). The colocalization of key small RNA components and a direct association with newly exported transcripts place P granules as a central compartment for RNA surveillance.

Adjacent to P granules are *Mutator* foci, which are considered hubs of small RNA amplification. *Mutator* foci are nucleated by the intrinsically disordered protein, MUT-16, which recruits key secondary siRNA synthesis proteins, including the RdRP RRF-1 and all *mutator* complex proteins (Phillips et al., 2012; Uebel et al., 2018). Recent work proposes that a *mutator* complex protein, MUT-2/RDE-3, a nucleotidyltransferase, marks transcripts for secondary siRNA synthesis by the addition of long 3’ poly-UG (pUG) repeats (Shukla et al., 2020). Loss of *Mutator* foci results in loss of many secondary siRNAs and an inability to respond to exogenous RNAi (Ketting et al., 1999; Phillips et al., 2012; Zhang et al., 2011). Therefore, *Mutator* foci represent a compartment of nuage in which siRNA amplification is organized.

A third nuage compartment, Z granules, are found between P granules and *Mutator* foci (Wan et al., 2018). Z granules are named after the first identified component, a conserved RNA helicase, ZNFX-1, which interacts with a number of small RNA proteins including the RdRP EGO- 1, and the Argonaute proteins CSR-1, WAGO-1, and PRG-1 (Ishidate et al., 2018). Notably, Z granules also contain WAGO-4, a secondary Argonaute required for transmission of secondary siRNAs to progeny; *znfx-1* mutants respond normally to RNAi, but are unable to pass the silencing signal on to progeny (Wan et al., 2018; Xu et al., 2018). Additionally, ZNFX-1, in coordination with other factors such as PID-2/ZSP-1 and LOTR-1, appears to balance the production of some secondary siRNAs across transcripts (Ishidate et al., 2018; Marnik et al., 2022; Placentino et al., 2021). Z granules, therefore, organize the transgenerational inheritance of small RNAs and balance the distribution of secondary siRNA synthesis across targeted transcripts.

The most recently discovered nuage compartment involved in siRNA pathways are SIMR foci, of which only three associated proteins are currently known: SIMR-1, an extended Tudor domain protein, HPO-40, a SIMR-1 paralog, and RSD-2 (Manage et al., 2020). SIMR-1 was first identified in a MUT-16 immunoprecipitation, but found to create spatially-separate foci adjacent to *Mutator* foci. The name SIMR-1 is derived from <u>si</u>RNA-defective and <u>m</u>ortal germline, as *simr-1* mutants lose some *mutator-*dependent siRNAs and become sterile over multiple generations at elevated temperature (Manage et al., 2020). Interestingly, *simr-1* mutants de-silence a piRNA sensor and lose small RNAs specifically mapping to piRNA targets (Manage et al., 2020). This mutant phenotype, along with a conserved role of Tudor domain proteins in both piRNA pathways and in protein-protein interactions (Pek et al., 2012), leads to the hypothesis that SIMR foci act in part to direct piRNA target genes to *Mutator* foci for downstream siRNA amplification (Manage et al., 2020). Furthermore, RSD-2 acts in the exogenous siRNA pathway to ensure efficient RNAi, suggesting that SIMR foci are a compartment of nuage that more generally mediates the transition between primary and secondary small RNA pathways (Han et al., 2008; Manage et al., 2020; Sakaguchi et al., 2014; Zhang et al., 2012).

All four of these compartments are not bound by lipid membranes. Specifically, P granules, Z granules, and *Mutator* foci have been shown to form via biomolecular phase separation, a process by which proteins and RNA de-mix from the surrounding bulk phase to form distinct, liquid-like condensates held together by transient, weak interactions and/or multivalent interactions. P granules exhibit controlled dissolution and condensation in embryos, rapid intramolecular rearrangement, and the ability to drip and fuse off nuclei when a shearing force is applied (Brangwynne et al., 2009). P granules also dissolve during heat stress and in 1,6- hexanediol, an aliphatic alcohol that disrupts the weak hydrophobic interactions that promote phase separation (Putnam et al., 2019; Updike et al., 2011). Z granules are colocalized with P granules until later in development, when they de-mix to form separate, adjacent foci, which also exhibit rapid molecular rearrangement (Wan et al., 2018). Z granule viscosity is regulated by the <u>p</u>iRNA-induced silencing <u>d</u>efective/<u>Z</u> granule <u>s</u>urface <u>p</u>rotein PID-2/ZSP-1 (Placentino et al., 2021; Wan et al., 2021). Finally, *Mutator* foci are concentration dependent, dissolve during heat stress and 1,6-hexanediol treatment, and are able undergo rapid molecular exchange with the surrounding bulk phase (Uebel et al., 2018). Thus, at least four distinct perinuclear nuage compartments, multiple of which are phase separated, are involved in RNA surveillance.

The initial characterization of these compartments demonstrates individual contributions to small RNA pathway organization, but the physical interaction and spatial configuration of all four compartments has only been briefly explored. It is also unclear how biomolecule exchange of RNAs, proteins, or small RNAs occurs between compartment boundaries. The current incomplete understanding of the *C. elegans* nuage assemblage limits formation of a comprehensive model to describe how multiple phase-separated compartments organize small RNA pathways and facilitate RNA silencing. Here we demonstrate that *Mutator* foci and P granule separation is maintained in ectopic environments and that interaction between compartments is dynamic and able to be re-established after perturbation. We use 3D-Structured Illumination Microscopy (SIM) to visualize the spatial organization of *Mutator* foci, P granules, Z granules, and SIMR foci. We discover a previously uncharacterized toroidal morphology of P granules, which we term “P granule pockets”, that interacts with all currently known compartments of nuage. Moreover, we quantify the relationships between germ granule compartments to reveal discrete populations of nuage which appear to assemble in a hierarchical manner. Finally, our quantification of RNA interaction with germ granule compartments demonstrates that nuage populations associated with *Mutator* foci are more likely to interact with RNAi-targeted RNAs. Ultimately, our data constructs a more cohesive model of how multiple phase-separated condensates organize small RNA pathways.

## RESULTS

### Granule separation is maintained in an ectopic environment

In the endogenous germline environment, *Mutator* foci and P granules exist as separate, yet adjacent, phase-separated perinuclear condensates (Phillips et al., 2012). We were interested in determining how these two compartments with liquid-like properties exist adjacently at the nuclear periphery without co-mixing. We first visualized the juxtaposition between *Mutator* foci and P granules in the germline at widefield resolution using endogenously expressed MUT-16::mCherry and PGL-1::GFP, respectively (Figure 1A). To determine if individual proteins within P granules or *Mutator* foci are sufficient to prevent co-mixing between compartments, we overexpressed MUT-16 and PGL-1 in *C. elegans* muscle tissue using the *myo-3* promoter (Figure 1B). MUT-16 is required for *Mutator* foci formation and has been previously shown to form ectopic foci in muscle cells when overexpressed, and PGL-1 is a major constituent of P granules that forms ectopic foci when overexpressed in intestines (Uebel et al., 2018; Updike et al., 2011). In previous work, we determined that *myo-3*-driven overexpression of either mCherry or GFP alone did not form condensates, thus overexpression of MUT-16 and PGL-1 is driving ectopic condensate formation (Uebel et al., 2018). Similar to P granule and Mutator foci interaction at the germline nuclear periphery, the PGL-1::GFP and MUT-16::mCherry condensates maintain their separate and adjacent relationship in the ectopic muscle environment (Figure 1B). To ensure that the ectopic interaction is not tag-specific, we also overexpressed PGL-1::mKate2 and MUT-16::GFP in muscle cells and obtained the same result (Figure S1A). Thus, the separate and adjacent relationship between P granules and *Mutator* foci does not require germline cellular environment or association with nuclear pores, but instead relies on the intermolecular interactions or phase-separation properties of key proteins in each condensate.

**Figure 1.**
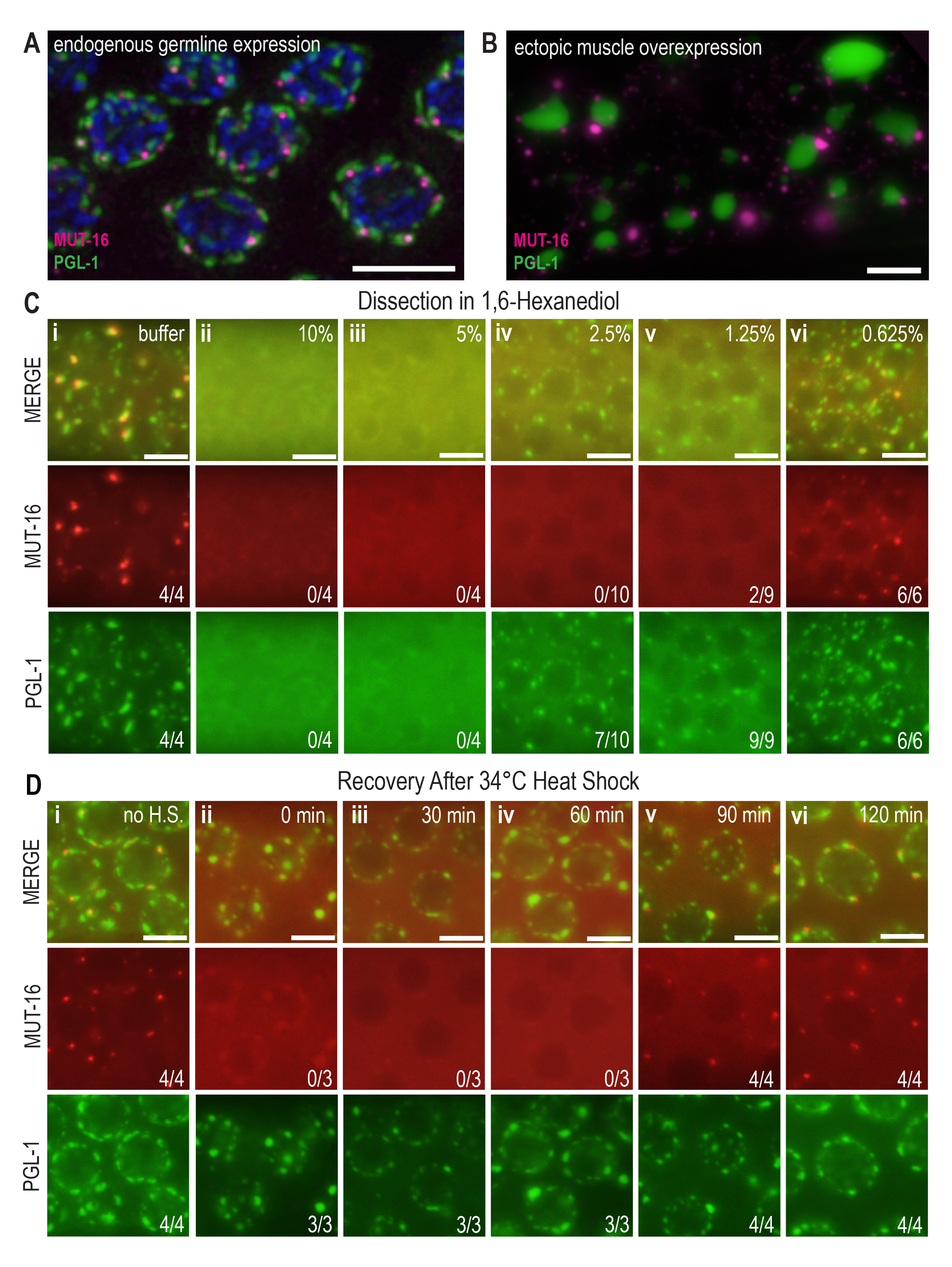
*Mutator* foci and P granule separation is independent of nuclear association and can be reestablished after perturbation. (A) Widefield immunofluorescence of *mut-16::mCherry; pgl-1::gfp* germlines shows that endogenous *Mutator* foci (MUT-16, magenta), and P granules (PGL-1, green), are adjacent yet distinct compartments. (B) Ectopically expressed MUT-16::mCherry (magenta) and PGL-1::GFP (green) driven by the *myo-3* muscle-specific promoter create condensates that maintain separation in the muscle environment. (C) Top: Live images of the transition zone of *mut-16::mCherry; pgl-1::gfp* gonads dissected in M9 buffer (Ci) or varying concentrations of 1,6- hexanediol (Cii-vi). Middle: *Mutator* foci (MUT-16, red) are dispersed in all concentrations of 1,6- hexanediol except 0.625%. Numbers indicate how many gonads displayed *Mutator* foci out of total assessed gonads. Bottom: P granules (PGL-1, green) are disrupted in only 10% and 5% 1,6- hexanediol, indicating different sensitivities to perturbation of weak hydrophobic interaction s. Numbers indicate how many gonads displayed P granules out of total assessed gonads. (D) *mut-16::mCherry; pgl-1::gfp* animals were subjected to heat stress at 34 °C for 2 hours and allowed to recover at room temperature (∼21 °C) for 2 hours. Top: Representative live images of the pachytene region were collected before heat stress (Di, no h.s.), immediately after heat stress (Dii, 0 min recovery), and for every 30 minutes during recovery (Diii-Dvi). Middle: *Mutator* foci (MUT-16, red) weakly colocalizes with P granules immediately after heat shock. At 30- and 60- minutes room temperature recovery, *Mutator* foci loses colocalization with P granules but remains dispersed in the cytoplasm. By 90- and 120- minutes room temperature recovery, MUT-16 reappears as separate, punctate foci adjacent to P granules, indicating the interaction is able to be re-established after perturbation. Numbers indicate gonads displaying punctate *Mutator* foci out of total assessed gonads. For Dii, colocalization with PGL-1 was seen in 3/3 gonads. Bottom: P granules do not completely disperse after 2 hours 34 °C heat stress. Numbers indicate how many gonads displayed P granules out of total assessed gonads. Scale bars, 5 µm.

To test if the molecular interactions contributing to phase separation differ significantly between *Mutator* foci and P granules, we subjected *mut-16::mCherry; pgl-1::gfp* gonads to 1,6- hexanediol, an aliphatic alcohol which disrupts weak, hydrophobic interactions and dissolves phase-separated condensates (Kroschwald et al., 2015). In a buffer control dissection, P granules and *Mutator* foci are clearly visible (Figure 1Ci). Both P granules and *Mutator* foci dissolve in 10% hexanediol and 5% hexanediol, as previously observed (Figure 1Cii, Ciii) (Uebel et al., 2018; Updike et al., 2011). We further dilute the concentration of hexanediol and observe that P granules are present at both 2.5% and 1.25% hexanediol, yet *Mutator* foci remain dissolved (Figure 1C iv, v). *Mutator* foci are present only in very dilute 0.625% hexanediol (Figure 1Cvi). Repeating the experiment with GFP-tagged *Mutator* foci yields similar results (Figure S1B). These data indicate that *Mutator* foci are more sensitive to perturbations of hydrophobic interactions via hexanediol than P granules, and this differential reliance on hydrophobicity for phase separation may be one factor that prevents co-mixing between adjacent condensates.

### P granule and *Mutator* foci interaction is dynamic

Many phase-separated condensates, including *Mutator* foci, dissolve with the addition of heat (Nott et al., 2015; Uebel et al., 2018). We previously discovered that *Mutator* foci dissolved by heat stress quickly reassemble as punctate foci during room temperature recovery (Uebel et al., 2018). To determine if *Mutator* foci re-establish adjacency to P granules after heat-stress dissolution, we subjected *mut-16::mCherry, pgl-1::gfp* animals to heat shock for 2 hours at 34 °C. Upon immediate imaging after removal from heat, we discovered that MUT-16 no longer forms punctate *Mutator* foci, but instead faintly colocalizes with P granules in enlarged foci (Figure 1Dii). In previous MUT-16::GFP heat-stress experiments, we observed a similarly enlarged *Mutator* focus phenotype, but did not have P granule markers to compare localization (Uebel et al., 2018). The two-hour heat shock did not completely dissolve P granules, but many detach from the nuclear membrane and collect in the syncytial gonad core (Figure S1C). Incomplete dissolution is likely an intermediate phenotype, as previous work demonstrates increased dissolution of PGL-1 after 3 hours at 34 °C heat stress (Jud et al., 2008; Uebel and Phillips, 2019). By 30- and 60- minutes room temperature recovery post heat shock, MUT-16 is less visibly colocalized with P granules and MUT-16 cytoplasmic signal remains high (Figure 1Diii-iv). By 90- and 120-minutes recovery post heat shock, *Mutator* foci appear strikingly similar to the non-heat-shocked control and are punctate and adjacent to P granules (Figure 1Di,1Dv-vi). The ability to quickly re-establish granule interaction after disruption suggests the adjacent relationship between P granules and *Mutator* foci is not necessarily established at an earlier developmental time point and continually maintained, but rather that *Mutator* foci may disassemble and reassemble next to particular P granules.

### Some P granules exhibit a toroidal morphology

To facilitate RNAi, exchange of biomolecules must occur within *C. elegans* nuage, but the physical interface between the different phase-separated compartments is not well understood. We sought to gain a high-resolution view of the interface between *Mutator* foci and P granules using 3D-Structured Illumination Microscopy (3D-SIM), which has nearly two-fold higher resolution than widefield microscopy (Gustafsson et al., 2008). In the transition zone, 3D-SIM shows *Mutator* foci and P granules interact adjacently as previously observed in widefield microscopy (Figure 2A). Unexpectedly however, imaging the mid-and late-pachytene regions revealed a previously uncharacterized P granule morphology; In these pachytene regions, a subset of P granules appear to surround *Mutator* foci in a toroidal “donut-like” morphology, which we further refer to as P granule pockets (Figure 2B-C, arrow, insets). We also observe a gap between *Mutator* foci and the encircling P granule, suggesting the two condensates do not directly interact in P granule pockets (Figure 2C insets), consistent with previous ideas of compartment interaction (Manage et al., 2020; Wan et al., 2018). After identifying P granule pockets at high resolution, we began to recognize P granule pockets in widefield microscopy. P granule pockets are visible in widefield live-imaging of undissected gonads, though the gap between P granules and *Mutator* foci is not resolved (Figure S2A). Because P granule pockets are present in live samples, the unique morphology does not appear to be a result of gonad fixation. To test if fluorescent protein tags influence the formation of P granule pockets, we used anti-PGL-1 to visualize untagged P granules and used anti-HA to visualize *mut-16::SNAP::HA*, which contains a SNAP-tag approximately 10 kD smaller than an mCherry tag. Subsequent 3D-SIM imaging revealed that antibody-visualized untagged P granules still form P granule pockets, further validating the authenticity of the P granule structures (Figure S2B). Of note, Pitt et al. (2000) briefly describes arch-shaped P granules, which may be related to P granule pockets. Altogether, our high-resolution imaging reveals a previously uncharacterized toroidal P granule morphology, in which some P granules encircle *Mutator* foci in the mid-and late-pachytene regions.

**Figure 2.**
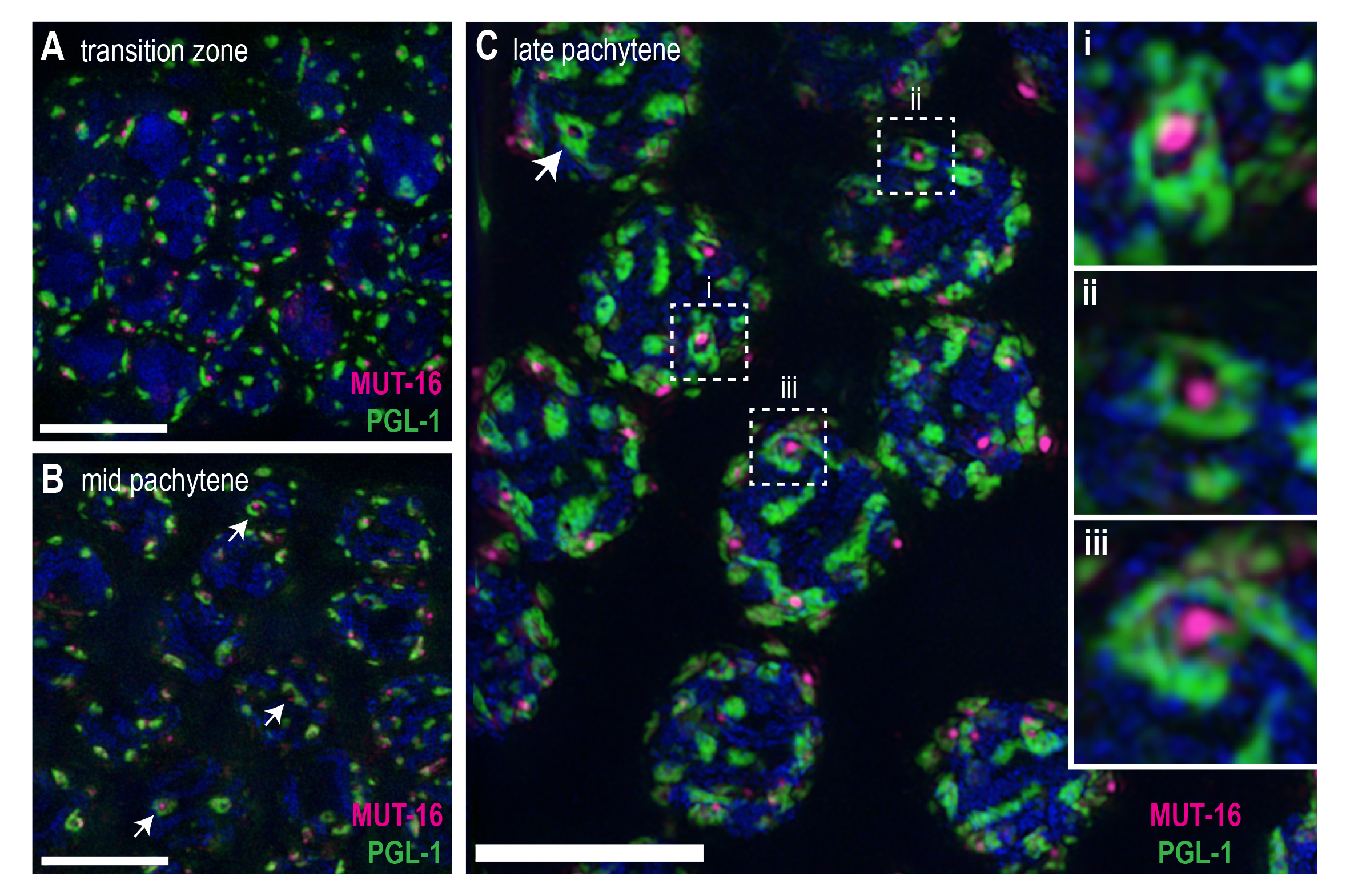
P granules form unique pocket morphologies in the mid and late pachytene. 3D-Structured Illumination Microscopy images of immunostained *mut-16::mCherry; pgl-1::gfp* germlines display P granules (PGL-1, green) and *Mutator* foci (MUT-16, magenta). (A) P granule and *Mutator* focus are adjacent to one another in the transition zone as previously described. (B) Some P granules appear to form an arc or circular morphology (arrows) around *Mutator* foci in the mid pachytene. (C) A toroidal, “donut-like” P granule morphology is readily apparent in the late pachytene (insets, arrow). Insets highlight the unique morphology, termed “P granule pocket”. Each P granule pocket appears to surround a *Mutator* focus, yet a gap is maintained between the foci. Images for (A) and (B) are comprised of 10 maximum projection Z-stacks (0.125 µm Z-step). Image (C) is comprised of 55 maximum-projection Z stacks (0.125 µm Z-step). Scale bars, 5 µm.

### Nuage compartments exist in distinct populations

From our microscopy, we observed that not all P granules are associated with a *Mutator* focus. We hypothesized that nuage compartments interact in distinct populations and sought to determine the stoichiometry between P granules, Z granules, *Mutator* foci, and SIMR foci to define nuage populations. To avoid possible artifacts from non-specific antibody binding, we used native fluorescence from endogenously tagged MUT-16, ZNFX-1, PGL-1 and SIMR-1 at widefield resolution. Quantification of fluorescent signal revealed that each nucleus (n = 30) in the late pachytene is associated with on average 22.4 (± 3.6) P granules (P), 18.4 (± 2.0) Z granules (Z), 12.4 (± 2.3) SIMR foci (S), and 10.8 (± 2.3) *Mutator* foci (M) (Figure 3A). P granules and Z granules are roughly twice as abundant as *Mutator* foci (2.1 and 1.7 fold, respectively). SIMR foci are more similar in quantity to *Mutator* foci, yet still have statistically significant higher foci counts per nucleus than *Mutator* foci (p = 0.007, Figure 3A). Our quantifications reveal that P granules are the most abundant nuage compartment in the germline, closely followed by Z granules, with *Mutator* foci as the scarcest nuage compartment.

**Figure 3.**
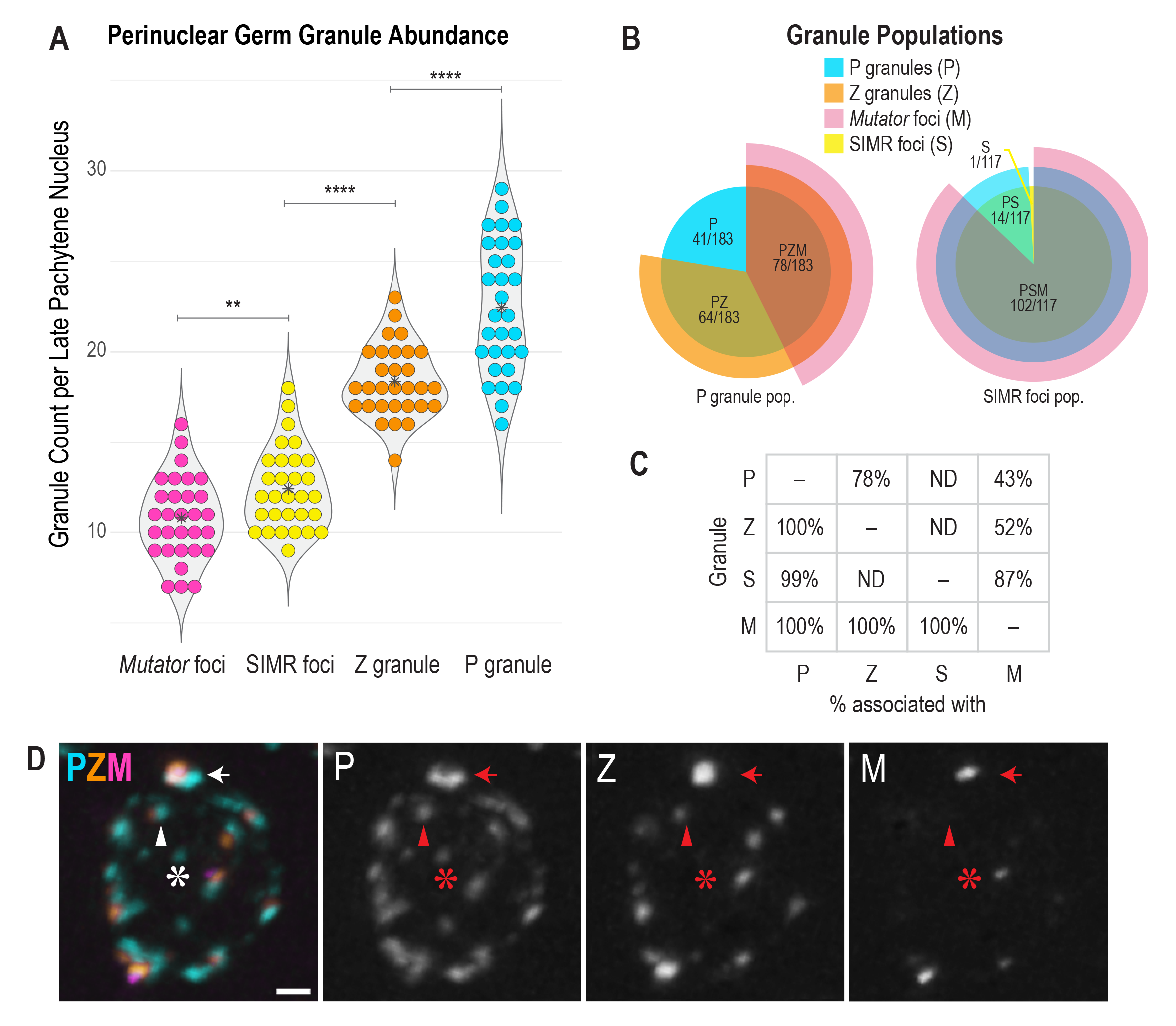
Nuage compartments exhibit a hierarchical stoichiometry. (A) Violin plot of fluorescently tagged germ granules surrounding nuclei in the late pachytene, with each dot corresponding to one nucleus (n = 30). Asterisks indicate average foci per nucleus. ** indicates p-value ≤ 0.01, **** indicates p-value ≤ 0.0001. Significance was determined with a two-tailed equal variance Student’s *t*-test. (B) Manual adjacency quantification from either *mut-16::gfp; rfp::znfx-1; pgl-1::bfp* to determine P granule populations (left) or *mut-16::gfp simr-1::mCherry; pgl-1::bfp* to determine SIMR foci populations (right). Overlapping pie charts reveal distinct populations of granule association. (C) Summary of combined granule stoichiometry indicating the percent that any one compartment (y-axis) is adjacent to a second compartment (x-axis). Of note, *Mutator* foci are always associated with all nuage compartments (PZM, n = 183 and PSM, n = 117), suggesting a hierarchical assembly of compartments. (D) Representative widefield image of a fixed *mut-16::gfp; rfp::znfx-1; pgl-1::bfp* late pachytene nucleus displaying the different P granule populations: P granule only (P, asterisk), P granule associated with Z granule (PZ, arrowhead), P granule associated with both Z granule and *Mutator* focus (PZM, arrow). Scale bar, 1 µm.

While our quantification demonstrates the abundance of each nuage compartment in a given nucleus, it is limited in directly addressing the relationships between the different compartments. To accurately assess compartment associations, we simultaneously visualized P granules, Z granules, and *Mutator* foci and manually evaluated each compartment for proximity to the other nuage compartments. We found three main populations of P granules (n = 183) (Figure 3B-D). The first population consists of P granules unassociated with any other visible compartment (P, 22%) (Figure 3B-C). These solitary P granules are generally smaller than other P granules (Figure 3D, asterisk and S3A). The second population contains P granules associated only with Z granules (PZ, 35%) (Figure 3B-C). This population ranges more broadly in size, but is generally comprised of medium and large P granules (Figure 3D, arrowhead and S3A). The third and most prevalent population, constituting 43% of assessed P granules, associates with both Z granules and *Mutator* foci (PZM) (Figure 3B-C). The majority of these P granules are large and include P granule pocket conformations (Figure 3D, arrow and S3A). 100% of assessed Z granules and *Mutator* foci were adjacent to P granules, indicating there are no solitary Z granules or *Mutator* foci (Figure 3C). To incorporate all known nuage compartments into our analysis, we next assessed the proximity of SIMR foci (S) to P granules and *Mutator* foci, and found that the majority of SIMR foci are adjacent to both compartments (PSM, 87%) (Figure 3B, S3B). In corroboration with our nuage compartment quantification, some SIMR foci were not associated with *Mutator* foci (PS, 12%). Only one SIMR focus did not appear to be associated with any other granule (Figure 3B-C), however Manage et al. (2020) finds that 100% of SIMR foci are adjacent to P granules and all SIMR foci are closely associated with Z granules (100%). Lastly, we find that all *Mutator* foci are associated with SIMR foci (100%) (Figure 3C, Figure S3B). From these relationship ratios, we extrapolate that P granules associated with *Mutator* foci are also associated with all other known germ granules and that PZSM constitutes 43% of all P granule populations. Moreover, nuage compartment relationships suggest a hierarchy of nucleation, in which P granules constitute a base granule for the subsequent nucleation of Z granules, SIMR foci, and *Mutator* foci, respectively.

### All nuage compartments are arranged within P granule pockets

Because not all P granules form P granule pockets, we quantified P granule pockets per nucleus visible at widefield resolution and discovered that nuclei in the late pachytene (n = 18) have on average 3.8 (±1.2) P granule pockets (Figure S3C). Each P granule pocket (n = 39) asso ciates with a *Mutator* focus. Therefore, all nuage compartments are present in P granule pockets. To assess the physical interaction between P granule pockets and Z granules or SIMR foci, we used SIM to visualize RFP::ZNFX-1, SIMR-1::GFP, and P granules. SIMR focus size was variable, but we were often able to detect a gap between SIMR-1 and P granules (Figure 4A, inset), indicating their localization within P granule pockets is similar to *Mutator* foci. In contrast, we found that Z granules generally occupy the entirety of P granule pocket interiors (Figure 4A), and can also appear to encompass *Mutator* foci in pocket-like formations (Figure 4B-C, inset). This experiment suggests that P granules directly interact with Z granules, and that Z granules bridge the gap between P granule pockets and *Mutator* foci and SIMR foci, similar to previous descriptions of PZM interaction (Manage et al., 2020; Wan et al., 2018).

**Figure 4.**
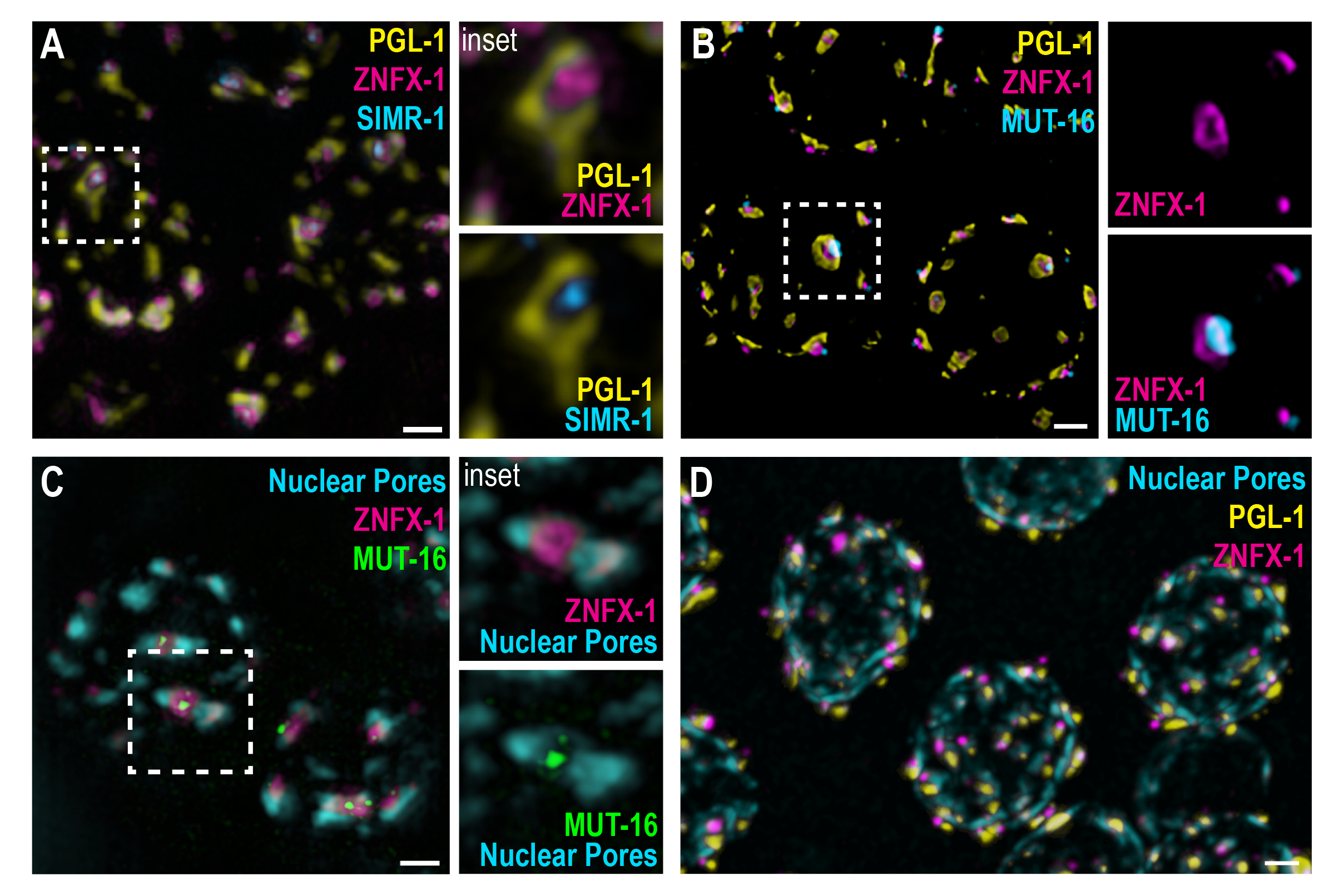
P granule pockets exhibit an exterior-to-interior organization. (A) Structured illumination of immunostained germlines with endogenously tagged SIMR-1 to detect SIMR foci (cyan), and ZNFX-1 to detect Z granules (magenta). P granules (yellow) are visualized with anti-PGL-1. Inset: a Z granule occupies the entire interior of the P granule pocket and a SIMR focus is innermost still. (B) Confocal image of fixed pachytene nuclei from a *mut-16::gfp*; *tagRFP::znfx-1*; *pgl-1::bfp* germline. Inset: a Z granule (magenta) encompasses a *Mutator* focus (cyan). (C) Structured illumination of immunostained germlines with endogenously tagged ZNFX-1 (magenta) and MUT-16 (green). Nuclear pore complexes (cyan) are visualized with anti-Nup 107 (mAb414). Inset: a Z granule occupies the interior of the nuclear pore pocket and a *Mutator* focus localizes to the center. (D) Confocal image of the late pachytene region of a fixed *gfp::znfx-1*, *tagRFP::npp-9; pgl-1::bfp* germline demonstrates nuclear pore interaction with germ granules. Scale bars, 1 µm.

To investigate the trajectory of RNA through nuage, we sought to determine which compartments interact with nuclear pores and, therefore, with newly exported RNA. Pitt et al. (2000) reports that 75% of nuclear pores are adjacent to P granules. We immunostained nuclear pore complexes, Z granules, and *Mutator* foci and discovered that nuclear pore complexes create similar patterns as P granules, and as such, some nuclear pores are arranged in a similar morphology to P granule pockets, with rafts of pores surrounding the other germ granule compartments (Figure 4C). Z granules occupy the inner space of the nuclear pore pocket and appear to have minimal overlap with nuclear pores (Figure 4C, inset). *Mutator* foci are innermost still, and, similar to their position within P granule pockets, a distinct gap exists between *Mutator* foci and the surrounding nuclear pore complexes (Figure 4C, inset). We conclude that *Mutator* foci do not interact with nuclear pores and thus are not directly involved in the capture of newly exported RNAs. The consistent exterior-to-interior organization of P granule pockets, wherein P granules appear to be the predominant nuage compartment interacting with newly exported RNAs, suggests that RNA follows a distinct trajectory through the nuage compartments – P granules, to Z granules, to SIMR and *Mutator* foci.

We next sought to address whether nuclear pore association differs among nuage populations. By visualizing nuclear pore complexes, P granules and Z granules, we found that all perinuclear P granules, regardless of size or compartment interactions, appear to be associated with nuclear pores, while some nuclear pores could be seen that were not associated with P granules (Figure 4D), consistent with Pitt et al. (2000) that reports 25% of nuclear pores are unassociated with P granules. These data indicate that all germ granule configurations are in contact with nuclear pores and able to capture nascent RNAs.

### RNAi-targeted RNAs interact with multiple germ granule populations

To investigate whether RNAs targeted by RNA interference preferentially associate with germ granules of a particular composition, we performed RNAi against the germline-expressed gene *mex-6* and visualized association of *mex-6* RNA with P or PM granules by single-molecule fluorescence *in-situ* hybridization (SM-FISH). We observed that the amount of cytoplasmic *mex-6* RNA was substantially reduced in the germline of *mex-6* RNAi-treated animals, compared to animals on control RNAi for 6 hours (Figure 5A-B). In contrast, a control RNA, *oma-1*, that is not targeted by RNAi is visible throughout the cytoplasm of both control and *mex-6* RNAi animals (Figure 5A-B). While there is no significant enrichment of *mex-6* RNA in germ granules of gonads treated with control RNAi (Figure 5C), the *mex-6* RNA remaining after *mex-6* RNAi treatment appears to be enriched in the germ granules of oocyte nuclei (Figure 5D), as previously observed (Ouyang and Seydoux, 2022), indicating that RNA localization with nuage is RNAi-targeting dependent. Moreover, we observe that the *mex-6* RNA signal often localizes between the *Mutator* foci and P granules (Figure 5D). While we do not have the resolution to definitively conclude to which granule the RNA associates, these data are consistent with the association of RNA with Z granules, which localize between P granules and Mutator foci (Wan et al., 2018). In further support, Ouyang and Seydoux (2022) have shown that ZNFX-1 is essential for accumulation of RNAi-targeted RNAs in germ granules.

**Figure 5.**
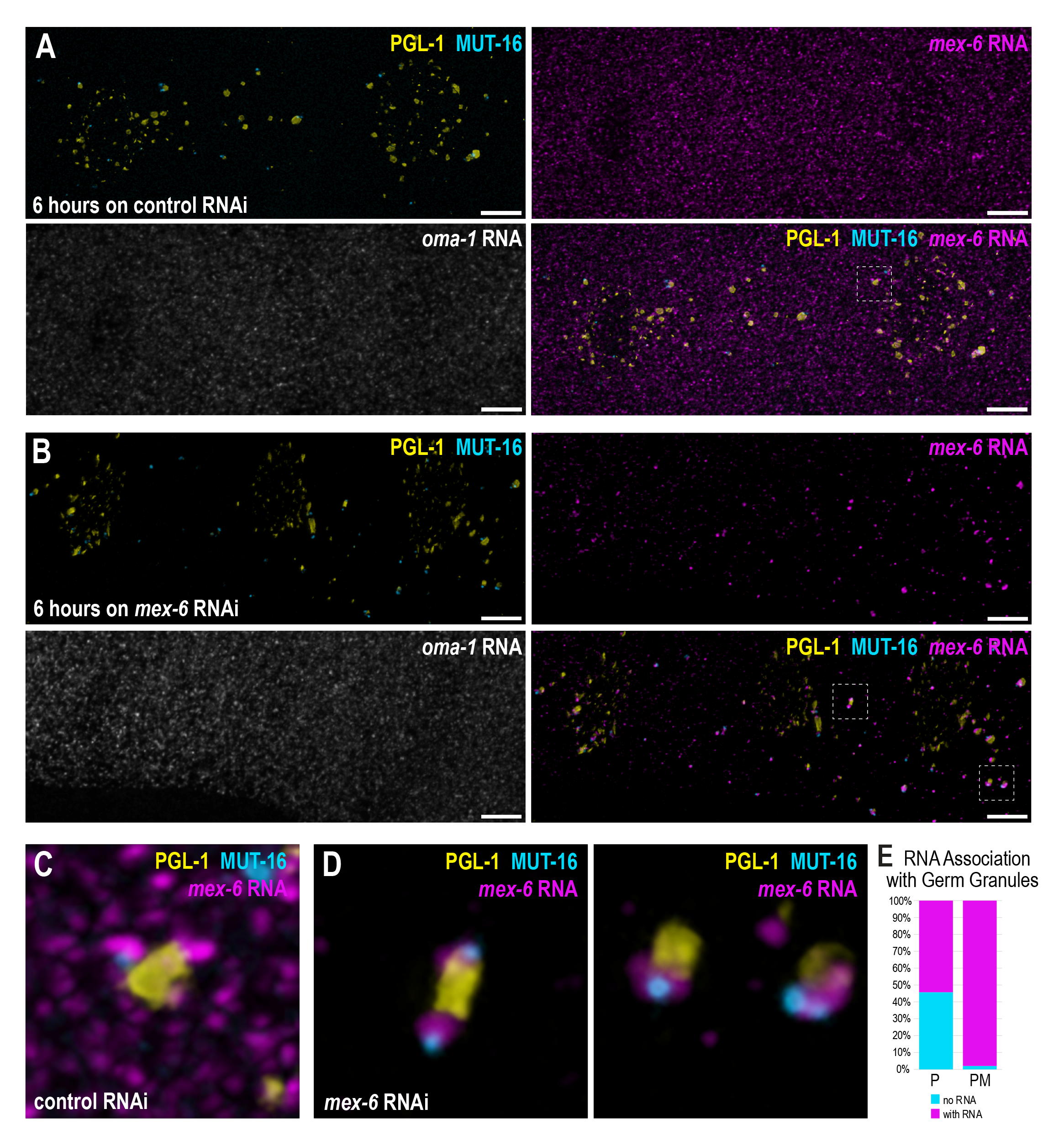
Silenced RNA associates preferentially with specific germ granule populations. (A-B) Confocal image of three oocytes in the diakinesis region of a *mut-16::gfp; pgl-1::bfp* germline following 6 hours on control (L4440) RNAi (A) or *mex-6* RNAi (B). Proximal oocytes are oriented to the right. (B) smFISH for both *oma-1* (control) and *mex-6* RNA shows that *mex-6* RNA, and not *oma-1* RNA, associates with germ granules after *mex-6* RNAi. Scale bars, 5 µm. (C-D) Insets from (A) and (B) showing *mex-6* RNA association with individual germ granules. For *mex-6* RNAi-treated animals, the *mex-6* RNA signal is located between the signals for PGL-1 and MUT-16. (E) Quantification of granules associated with RNA from *mut-16::gfp; pgl-1::bfp* following 6 hours on *mex-6* RNAi to determine frequency of P and PM interactions with *mex-6* RNA.

We next sought to determine the frequency with which *mex-6* RNA associates with P or PM granules. Note that when imaging only P granules and *Mutator* foci, we can score “P” (inclusive of P, PZ, and PZS granules) or “PM” (PZSM granules). We found that, of the granules scored as “P” (P, PZ, or PZS), 54% were associated with *mex-6* RNA while 46% were unassociated with RNA (Figure 5E). In contrast, the granules scored as “PM” (PZSM granules) were associated with *mex-6* RNA 98% of the time. These data indicate that germ granules that comprise all currently known subcompartments associate with RNA at a higher frequency than germ granules lacking *Mutator* foci (P, PZ, or PZS). Thus, while RNAi-targeted RNAs were not associated exclusively with one population of nuage, they also do not associate indiscriminately with the different nuage populations.

## DISCUSSION

### Physical organization of nuage

Previous to our work, *C. elegans* nuage had not been characterized using super-resolved microscopy techniques. Our work adds three key details to the physical organization of *C. elegans* nuage. First, nuclear pores do not appear to directly colocalize with Z granules or *Mutator* foci, and instead associate primarily with P granules (Figure 4). Supported by previous literature demonstrating a direct interaction between P granules and nuclear pores (Pitt et al., 2000; Sheth et al., 2010), we conclude that P granules are the first compartment to directly capture newly exported RNA (Figure 6). Second, a subset of P granules exhibit a toroidal morphology which encompasses all known nuage compartments in a consistent exterior-to-interior organization. This organization is similar to previous findings (Manage et al., 2020; Wan et al., 2018), but distinct in that the P granule surrounds a Z granule, which in turn encompasses both a SIMR focus and *Mutator* focus (Figure 6). Third, P granule size correlates with more complex arrangements of nuage compartments and the formation of P granule pockets (Figure S3). Our model of the spatial organization of nuage provides a foundation for understanding the trajectory of an RNA as it enters small RNA pathway compartments. In future studies, it will be necessary to determine if P granule pocket morphology and organization actively promotes RNA surveillance, or if it is simply a biophysical outcome of granule size and interaction.

**Figure 6.**
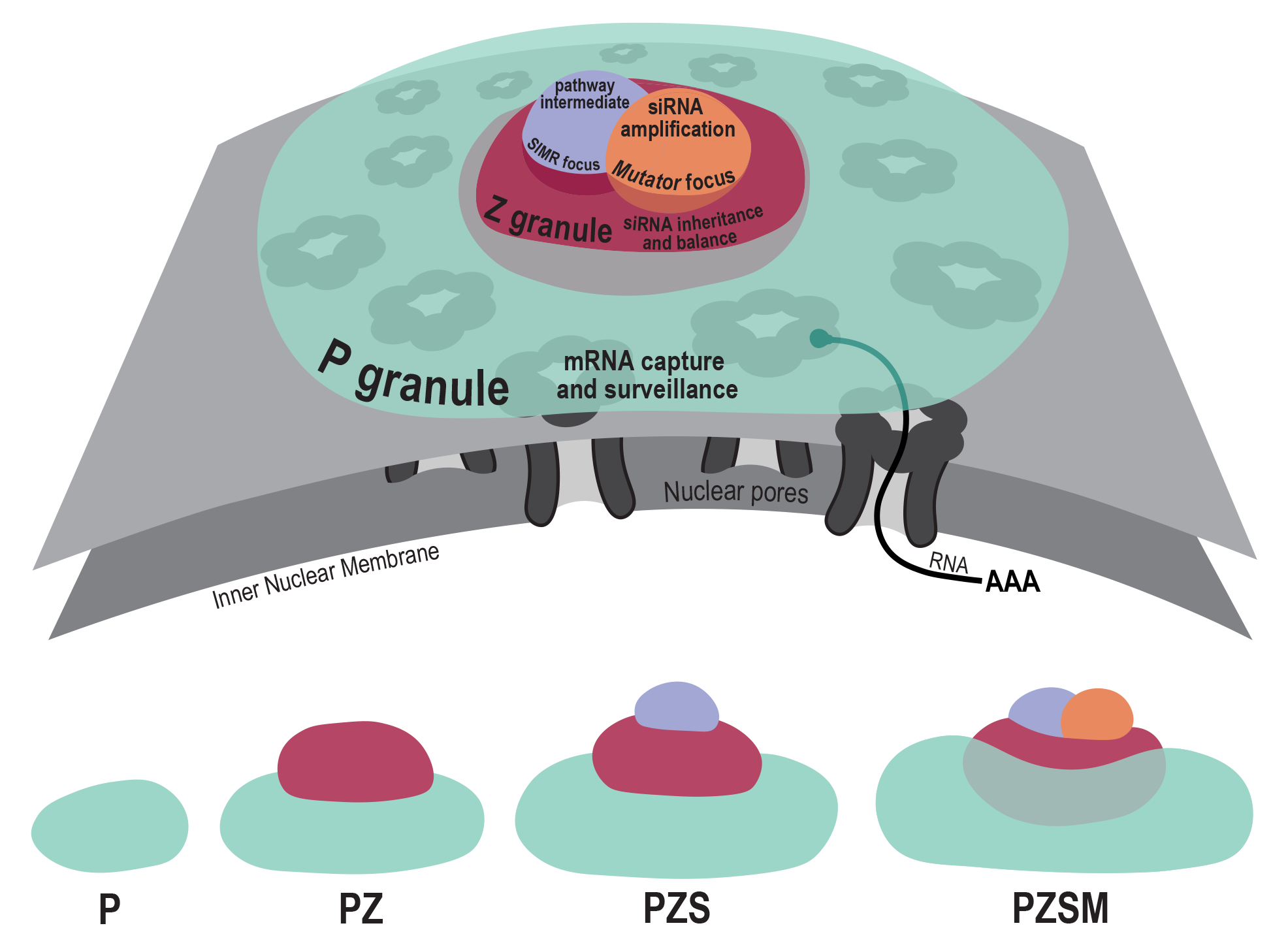
Working model of P granule pocket organization and nuage assembly. Top: Model of a P granule exhibiting toroidal “pocket” morphology at the periphery of a *C. elegans* germ cell nucleus (gray). Nuclear pores (dark gray) interact primarily with the P granule pocket (teal), enabling P granules to capture nascent RNA (black). The P granule pocket encircles a Z granule (red) which balances secondary siRNA synthesis across transcripts and is required for siRNA inheritance. The Z granule further encompasses a SIMR focus (purple), which acts as an intermediate between primary and secondary siRNA pathways, and a *Mutator* focus, which is required for secondary siRNA synthesis (orange). Bottom: Assembly hierarchy of P granule populations proceeding from P to PZSM (left to right). P granules associated with Mutator foci are associated with all known nuage compartments.

### Dynamics of germ granule interactions

We find that the separation of P granule and *Mutator* compartments does not rely on features of the germline environment, such as association with the nuclear periphery or germline mRNAs. Rather, separation of condensate-promoting proteins PGL-1 and MUT-16 can be maintained in an exogenous environment, as demonstrated by our ectopic overexpression experiments (Figure 1). Moreover, we find that PGL-1 and MUT-16 proteins have drastically different responses to perturbation of weak, hydrophobic interactions, suggesting that compartment separation may also be conferred by dissimilar compositions of the multivalent interactions that promote phase separation, such as differences in amino acid contents of intrinsically disordered regions. Finally, we show that following disruption, *Mutator* foci are able to re-form and restore adjacency to P granules. This has exciting implications for the dynamic nature of nuage compartments: *Mutator* foci may disassemble or reassemble adjacent to particular P granules, possibly in reaction to changing RNA and/or small RNA populations. Although this finding does not include the other nuage compartments, it begins to illuminate mechanisms that may maintain and promote multiple phase-separated structures as distinct compartments within nuage.

### Distinct compositions of germ granule populations

Our work reveals discrete populations of perinuclear nuage (P, PZ, PZS, PZSM) that appear to form in a hierarchical manner, in that the association of Z granules with P granules is required for the nucleation of SIMR foci and then *Mutator* foci (Figure 3). These results are consistent with prior work demonstrating that disruption of essential P granule components leads to disruption of Z granules and *Mutator* foci (Ishidate et al., 2018; Singh et al., 2021). Thus far, we cannot assign distinct functional roles for the different granule configurations in RNA surveillance, but our evidence that RNAi-targeted RNAs associate at a higher frequency with germ granules containing *Mutator* foci compared to germ granules lacking *Mutator* foci is encouraging (Figure 5). However, it is also possible that germ granule proteins and associated RNAs can freely exchange between germ granules via the cytoplasm. Thus, while germ granules may provide a hotspot for screening RNAs as they are exported from the nucleus, RNA surveillance need not be limited to granules with particular biochemical activities such as siRNA synthesis, or for that matter, to granules at all.

One caveat to this work is the possibility that germ granule compartments may be present below the level of detection of confocal microscopy or 3D-SIM. For example, PZS granules may contain *Mutator* complex proteins at visually undetectable but functionally relevant levels. Perhaps the compartments that we can visualize within a given germ granule are indicative of what molecular processes are occurring in that granule rather than indicating a restriction on what molecular processes can occur there. Ishidate et al. (2018) propose that subdomains of nuage could be formed by synchronous molecular events occurring on such a massive scale to become visible as distinct phase-separated domains within nuage. From this vantage point, we can speculate that the hierarchical assembly of germ granule compartments may be determined by the order of the molecular events required for RNA silencing. As we identify new RNA silencing factors associated with nuage and re-image previously characterized proteins at higher resolution, we may find more and more domains within nuage that define distinct compartments with unique molecular functions.

Together, our work uncovers new details on the organization and compartmentalization of *C. elegans* nuage and advances the understanding of how RNA surveillance is organized through multiple phase-separated compartments.

## MATERIALS AND METHODS

### C. elegans strains

| Strain # | Genotype | Source | Fig. |
| --- | --- | --- | --- |
| USC717 | mut-16(cmp3[mut-16::gfp::3xFLAG]) I | Uebel et al. 2018 | S1B |
| USC1242 | unc-119(ed3) III; cmpEx95[myo-3p::mut-16::gfp; myo3p::pgl-1::mKate] | this work | S1A |
| USC1266 | mut-16(cmp41[mut-16::mCherry::2xHA]) I; pgl-1(sam33[pgl-1::gfp::3xFLAG]) IV | Uebel and Phillips 2020 | 1A, 1C-D, 2A-C, S1C, S2A |
| USC1315 | mut-16(cmp249[SNAP::HA::mut-16]) I | this work | S2B |
| USC1353 | mut-16(cmp259 [2xMYC::bfp::mut-16]) simr-1(cmp112[simr-1::GFP::3xFLAG]) I; znfx-1(gg634[HA::tagRFP::znfx-1]) II | this work | 4A |
| USC1392 | unc-119(ed3) III; cmpEx97[myo-3p::mut-16::mCherry; myo-3p::pgl-1::gfp] | this work | 1B |
| USC1401 | simr-1(cmp15[simr-1::mCherry::HA]) mut-16(cmp3[mut-16::gfp::3xFLAG]) I; pgl-1(cmp226[pgl-1::bfp::3xFLAG]) IV | Uebel et al. 2021 | 3A-C, S3B |
| USC1408 | mut-16(cmp3[mut-16::gfp::3xFLAG]) I; pgl-1(cmp226[pgl-1::bfp 3xFLAG]) IV | this work | 5A-E |
| USC1409 | mut-16(cmp3[mut-16::gfp::3xFLAG]) I; znfx-1(gg634[HA::tagRFP::znfx-1]) II; pgl-1(cmp226[pgl-1::bfp::3xFLAG]) IV | this work | 3A-D, 4B, S3A, S3C |
| USC1410 | mut-16(cmp3[mut-16::gfp::3xFLAG]) I; znfx-1(gg634[HA::tagRFP::znfx-1]) II | this work | 4C |
| USC1532 | znfx-1(cmp294[GFP::3xFLAG::znfx-1]) II; npp-9 (gg654[tagRFP::3xFLAG::npp-9]) III; pgl-1 (cmp226[pgl-1::bfp::3xFLAG]) IV | this work | 4D |

### Primers

List of primers used to construct the strains used in this work.

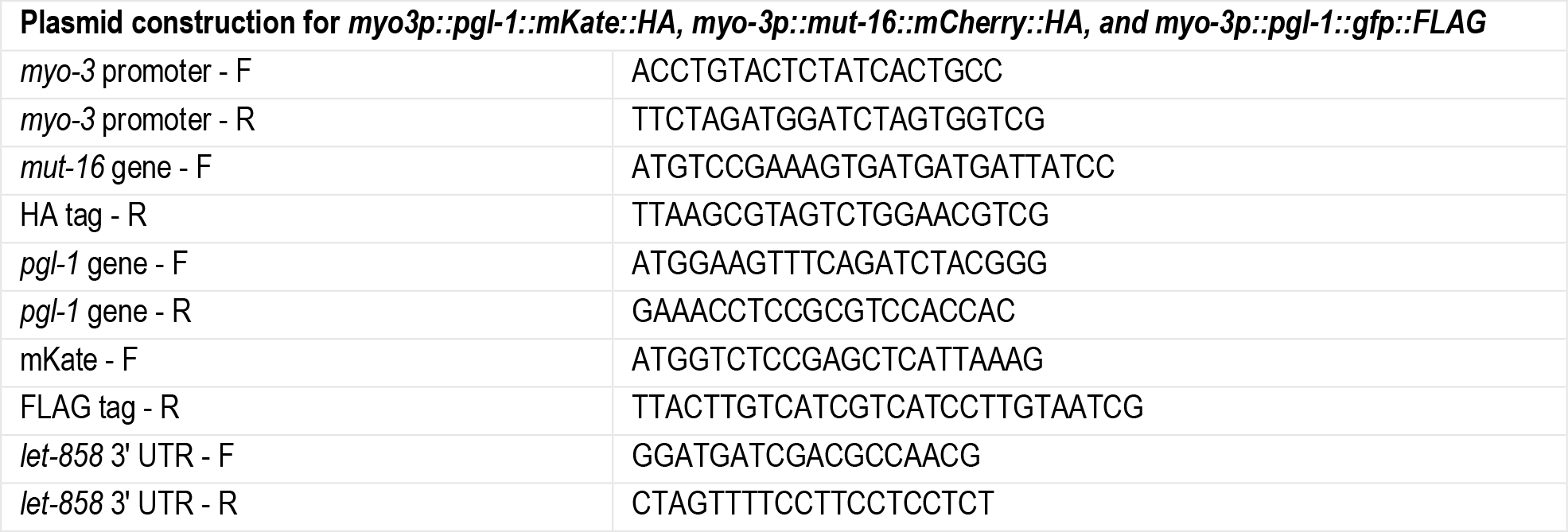
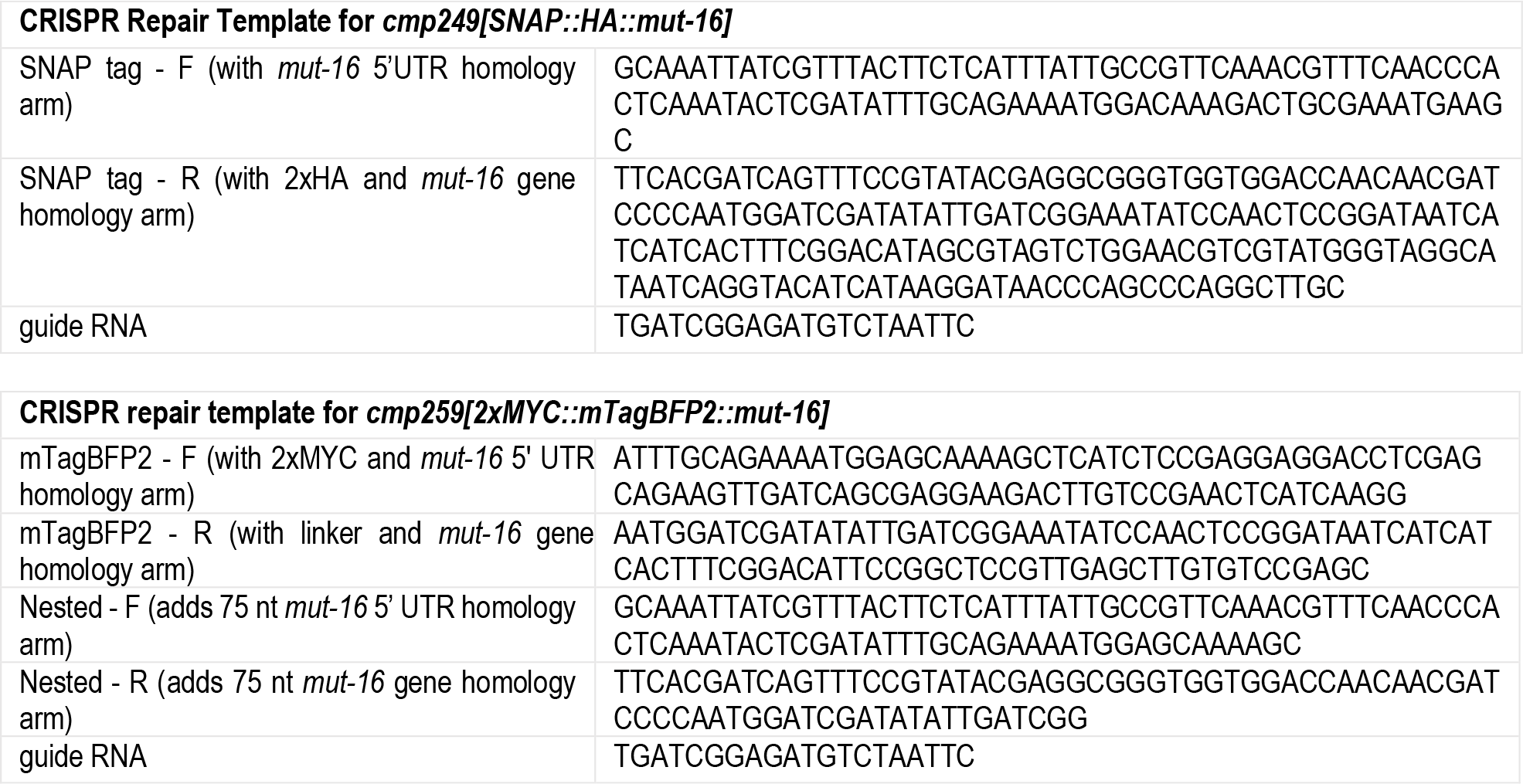
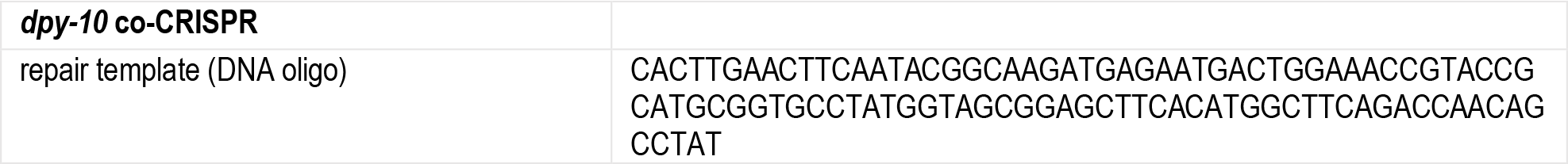

### Strain Construction

Ectopic expression plasmids for *myo3p::pgl-1::mKate::HA, myo-3p::mut-16::mCherry::HA, and myo-3p::pgl-1::gfp::FLAG*, were created via PCR amplification (see primer list) and assembled into a Spe-1 digested pCFJ151 (Addgene #19330) vector using isothermal assembly (Gibson et al., 2009). DNA templates for PCR products were as follows - *myo-3p* from pCFJ104 (Addgene #19328), *pgl-1::GFP::FLAG* from DUP75 (Andralojc et al., 2017), *mut-16::mCherry::HA* from USC896 (Uebel et al., 2018), *pgl-1* gene from N2 genomic DNA, and *let-858* 3’ UTR and *mKate::HA::let-858* 3’UTR from pDD287 (Addgene #70685). Correct sequences of constructed plasmids were confirmed with Sanger sequencing. *myo-3p::mut-16::gfp* was constructed previously (Uebel et al., 2018). We generated extra-chromosomal arrays as follows: 10 ng/µl *myo-3p::mut-16::gfp*, 10 ng/µl *myo-3p::pgl-1::mKate2* and 70 ng/µl pBluescript was injected into HT1593 *unc-119(ed3) III* and Unc-rescued animals were selected to create USC1242 (*cmpEx95*). 10 ng/µl *myo3p::mut-16::mCherry::2xHA*, 10 ng/µl *myo3p::pgl-1::GFP::3xFLAG*, and 70 ng/µl pBluescript was injected into HT1593 *unc-119(ed3) III* and Unc-rescued animals were selected to create USC1392 (*cmpEx97)*.

For *mut-16(cmp249[SNAP::HA::mut-16])* and *cmp259[2xMYC::mTagBFP2::mut-16]*, PCR repair templates for CRISPR genome editing were designed with primers (see primer list) and injected as previously described (Paix et al., 2017). Injection mix was prepared as follows: 2.5 µg/µL Cas9 protein (IDT, Cat# 1081059), 100 ng/µL tracrRNA (IDT, Cat# 1072534), 14 ng/µL *dpy-10* crRNA (IDT Alt-R system), 42 ng/µL gene-specific crRNA (IDT) were incubated at 37 °C for 10 minutes. Following incubation, 10 pmol of PCR repair template amplified from pSNAP-tag plasmid (Addgene #101135) or mTagBFP2 plasmid (Addgene #75029) and 110 ng/µL of *dpy-10* ssODN repair template were added. The injection mix was centrifuged at max speed, transferred to a fresh tube, and microinjected into N2 animals for *cmp249[SNAP::HA::mut-16* or into USC1229 (simr-1(cmp112[simr-1::GFP::3xFLAG]) I; znfx-1(gg634[HA::tagRFP::znfx-1])) for *cmp259[2xMYC::mTagBFP2::mut-16]*. F1 animals with Rol or Dpy phenotypes were singled to individual plates and screened for insertion of the tag by PCR. Transgenic animals were confirmed by Sanger sequencing.

All other new strains created for this study were made via mating available strains. Strains were maintained at 15 °C, 20 °C, or room temperature (∼21 °C) unless performing heat stress experiments.

#### Immunostaining

For fixed widefield, confocal, and 3D-SIM fluorescent imaging, adult animals were dissected in egg buffer containing 0.1% Tween-20 and fixed for 3 minutes in a 1% v/v formaldehyde solution. Animals were permeabilized via freeze-cracking and further fixed in ice-cold 100% methanol for 1 minute. Slide preparation was then performed as described in Phillips et al. (2009).

Antibody staining of fixed germlines was performed with 1:500 Rat anti-HA 3F10 (Roche 11867423001), 1:500 Rabbit anti-GFP (Thermo A-11122), 1:50 Mouse IgM anti-PGL-1 (DSHB K76), 1:2000 Mouse IgG anti-FLAG M2 (Sigma F-1804), and 1:5000 Mouse IgG anti-Nup107 (Covance mAb414). Fluorescent secondary antibodies used at 1:1000 include Goat anti-rabbit 488 (Thermo A-11008), Goat anti-mouse IgG 488 (Thermo A-11029), and Goat anti-rat 555 (Thermo A-21434). Secondary antibodies used at 1:500 include Goat anti-mouse IgM 647 (Thermo A-21238) and Goat anti-mouse IgG 647 (Thermo A21236).

### Live Imaging and Widefield Microscopy

Young adult animals were standardized for age by selection of the L4 larval stage on the day preceding imaging. For live imaging, undissected animals were mounted on glass slides in <1% Sodium Azide in M9 buffer solution to prevent movement. For dissected gonads, animals were quickly dissected in M9 with a #11 Feather Blade scalpel. Coverslip edges were sealed with clear nail polish to prevent buffer evaporation. All live imaging and some immunostaining were performed on a GE Healthcare DeltaVision Elite microscope using a 60x NA 1.42 oil-immersion objective. Unless otherwise stated, 5 Z-stacks (0.2 µm Z-step) were compiled as maximum intensity projections to create each image. Adobe Photoshop was used to pseudo-color and adjust the brightness/contrast of each image for clarity.

### Confocal Microscopy

Confocal imaging was performed on a Leica Stellaris 5 confocal microscope using a 63x NA 1.40 oil-immersion objective and the Lightening deconvolution package. Z-stack spacing was system optimized and maximum intensity projections of the entire nucleus or oocyte were generated using Fiji. Adobe Photoshop was used to pseudo-color and adjust the brightness/contrast of each image for clarity.

### 3D-Structured Illumination Microscopy

3D-SIM was performed at the USC Core Center of Excellence in Nano Imaging using a GE Healthcare DeltaVision OMX V4 microscope with an Olympus 60x NA 1.42 oil-immersion objective. 3 angles and 5 phases were collected for each Z-stack (0.125 Z-step). Laser wavelengths of 405 nm, 488 nm, 568 nm, and/or 642 nm were used for excitation. Image processing was performed with softWoRx software for structured illumination image reconstruction (Weiner filter = 0.005) and channel alignment. Final images were created by compiling maximum intensity projections of the reconstructions in softWoRx. Channels were pseudo-colored and brightness/contrast was adjusted for clarity in Adobe Photoshop.

### Granule Disruption

For 1,6-hexanediol experiments, young adult animals were dissected in either M9 buffer as a control, or a solution of 10%, 5%, 2.5%, 1.25%, or 0.625% 1,6-hexanediol in M9 buffer. Animals were imaged within 5 minutes after dissection. At least four gonads were assessed for each dilution.

For heat-shock experiments, NGM plates of young adult animals were wrapped in parafilm and placed in an incubator at 34 °C for 2 hours. For room temperature recovery, plates were allowed to recover on the benchtop at room temperature (∼21 °C) for the specified time. Images of undissected gonads were collected and imaged within ± 10 minutes from each indicated timepoint. At least 3 gonads were assessed for each time point.

### Single-Molecule Fluorescence *In Situ* Hybridization (smFISH)

48 smFISH probes were designed for each *mex-6* and *oma-1* mRNAs using the Stellaris Probe Designer, and labeled with Quasar570 and Quasar670 dyes, respectively. smFISH was performed on dissected germlines as described previously (Ouyang and Seydoux, 2022). Briefly, worms were dissected in 1x PBS and fixed using 3.7% (vol./vol.) EM-grade paraformaldehyde in 1X PBS for 15 minutes. Post-fixation, excess liquid was removed and animals were permeabilized via freeze-cracking, rinsed twice in 1x PBST and stored in 70% ice-cold ethanol overnight. Samples were washed with Stellaris Wash Buffer A solution (2 mL Stellaris RNA FISH Wash Buffer A (Biosearch Technologies SMF-WA1-60), 7 mL Nuclease-free water, and 1 mL deionized formamide) followed by 100 μL of freshly prepared Hybridization Buffer (90 μL Stellaris RNA FISH Hybridization Buffer (Biosearch Technologies, SMF-HB1-10), 10 μL deionized formamide, and 1ul of each 12.5 μM RNA FISH probe) and incubated at 37 °C overnight. The samples were briefly rinsed again in 500 μL of Stellaris Wash Buffer A mixture, then incubated in in 500 μL of Stellaris Wash Buffer A mixture for 30 min at 37 °C, washed with 500 μL Stellaris RNA FISH Wash Buffer B (Biosearch Technologies, SMF-WB1-20), and incubated in 500 μL Stellaris RNA FISH Wash Buffer B for 5 min at room temperature. Lastly, the samples were mounted in Vectashield Mounting Medium and sealed under a coverslip with nail polish.

### Granule and RNA Adjacency Quantification

P granule, Z granule, and *Mutator* foci quantification was performed on USC1409. Animals were dissected in 1x PBS and fixed for 20 minutes by adding 1:1 EM-grade paraformaldehyde for a 4% final solution. Animals were permeabilized via freeze-cracking, rinsed twice in 1x PBST and stored in 70% ice-cold ethanol overnight. Gonads were mounted in NPG glycerol and sealed with nail polish. The endogenous fluorescence of each granule was used for quantification to avoid any off-target or background artifacts that could arise from immunostaining. Widefield fluorescence images were taken on the GE Deltavision Elite and deconvolved with softWoRx software. 10 complete nuclei from the late-pachytene region of 3 gonads were selected at random for quantification for a total n = 30. TIFs for individual channels of each nucleus were converted to 8-bit files, thresholded in FIJI 3D Object Counter, and manually inspected to ensure appropriate granule coverage through all Z stacks. SIMR foci quantification was performed on USC1401 in the same manner. Granule counts were obtained from Fiji 3D Object Counter and plotted via ggPlot.

Granule adjacency quantification was performed on 3 randomly selected late pachytene nuclei from each of the 3 previously assessed germlines in both USC1401 for SIMR populations and USC1409 for P granule populations (n = 9 nuclei per genotype). To aid in adjacency visualization, fluorescent signal from each channel was converted into a 3D object using the Fiji 3D-Viewer plugin. Fluorescent thresholding was the same as for granule quantification, the resampling factor was set to 1, and the display was set to “surface”. All three channels were layered into one 3D-viewer image for adjacency assessment and was cross-checked with the original TIF files. For USC1409, P granules (n = 183), Z granules (n = 142), and *Mutator* foci (n = 78) were assessed. Raw quantification data shows 142 Z granules were adjacent to P granules, and 78 *Mutator* foci were adjacent to both P granules and Z granules. There were no solitary Z granules or *Mutator* foci. Of all P granules assessed, 6 P granules were associated with 2 Z granules (PZZ, 3%), 3 P granules had 2 Z granules and 1 *Mutator* foci (PZZM, 2%) and 1 P granule had 2 Z granules and 2 *Mutator* foci (PZMZM, 1%). These outliers were grouped as PZ and PZM, respectively. For USC1401, SIMR foci (n = 117) and *Mutator* foci (n = 102) were assessed. 116 SIMR foci were adjacent to P granules, and 102 *Mutator* foci were adjacent to both SIMR foci and P granules. P granules were not individually counted for USC1401. P granules for both USC1401 and USC1409 were manually assessed from the TIF files for P granule pocket morphology and qualitative size.

RNA adjacency quantification was performed on USC1408 using the Fiji 3D Objects counter plugin. 16-bit TIFs of maximum intensity projections for individual channels of each oocyte were thresholded in Fiji 3D Object Counter and manually inspected to ensure appropriate granule coverage. The size filter minimum was set to 10 and map display was set to “objects”. Object maps for each channel (P granules, Mutator foci, and RNA) were merged and manually inspected for object colocalization/overlap. 3 oocytes from 3 different gonads were quantified (n = 9 total oocytes), providing a total of 597 P granules for assessment of adjacency to Mutator foci and/or RNA. P granules were scored as solitary P granules (P = 208), P granules with RNA (PR =247), P granules with Mutator foci (PM = 3), or P granules with Mutator foci and RNA (PMR = 139).

## ACKNOWLEDGEMENTS

We would like to thank Kevin Keomanee-Dizon at the USC Core Center of Excellence in Nano Imaging for his expertise and training with 3D-SIM microscopy, and Scott Kennedy and Heng-Chi Lee for sharing strains.

## COMPETING INTERESTS

The authors declare no competing or financial interests.

## FUNDING

C.J.U. is funded by the National Science Foundation Graduate Research Fellowship Program (DGE 1418060), the USC Research Enhancement Fellowship, and the Chemical Biology Interface T32-GM118289 NIH/Dornsife grant. The Phillips lab is supported by R35 GM119656, and by the Pew Charitable Trusts (www.pewtrusts.org).

---

**Figure S1.**
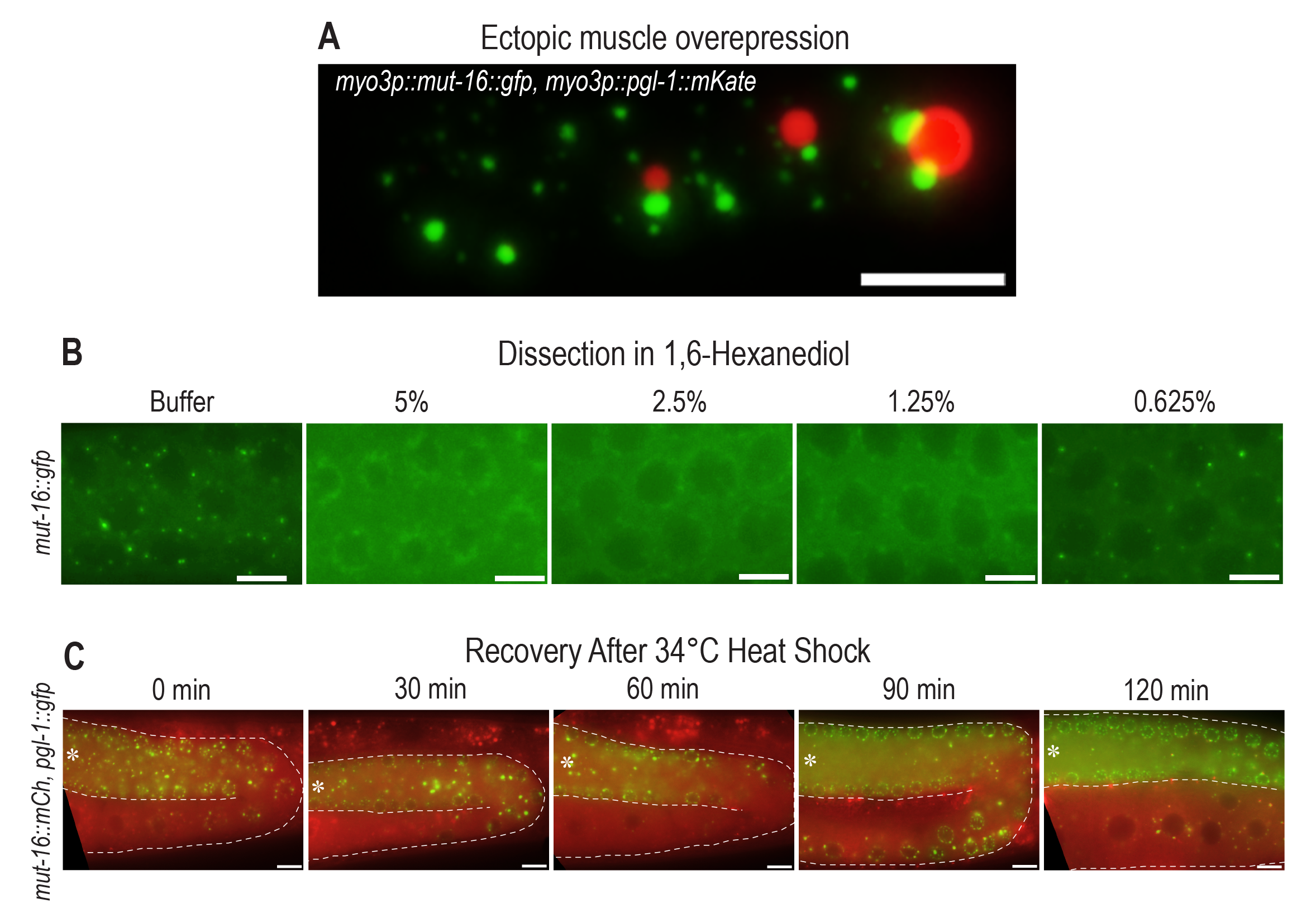
Ectopic interaction and dissolution of granules is consistent across multiple fluorescent tags. (A) Ectopically expressed MUT-16::GFP and PGL-1::mKate driven by the *myo-3* muscle specific promoter form distinct condensates that maintain separation in the muscle. (B) Live images of gonads dissected in M9 buffer or a series of 1,6-hexanediol dilutions reveals punctate MUT-16::GFP foci are present only in buffer and the most dilute 1,6-hexanediol concentration (0.625%). (C) Live images of *mut-16::mCherry; pgl-1::gfp* animals subjected to 2 hours heat shock at 34° C and allowed to recover at room temperature (∼21 °C) for the indicated times. Immediately after heat stress (0 min recovery) and following 30- and 60-minute recovery, P granules (green) are detached from the nuclear periphery and collect in the rachis. Fewer detached P granules appear in the rachis of animals after 90- and 120- minutes of room temperature recovery. Distal gonad region is oriented to the left (asterisk) with the proximal oocytes below. Scale bars, 5 µm.

**Figure S2.**
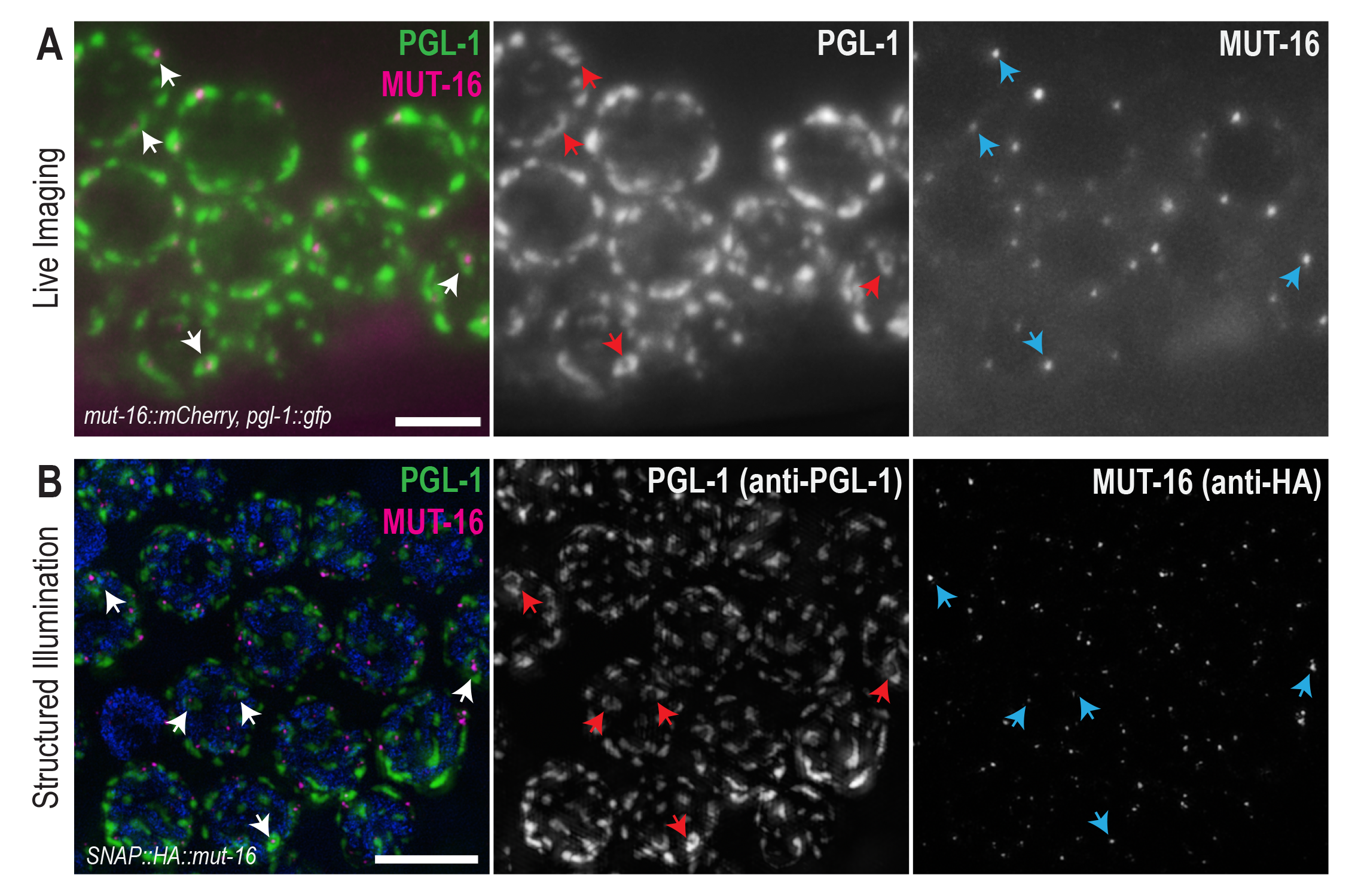
P granule pocket morphology is visible in widefield live imaging and in untagged P granules. (A) P granule pockets (red arrows) are detectable in widefield live imaging of the late pachytene regions of endogenously tagged *mut-16::mCherry; pgl-1::gfp* animals. The gap between *Mutator* foci (magenta) and P granules (green), which was visible in SIM imaging, is not apparent in widefield imaging (white arrows). Each P granule pocket is associated with a *Mutator* focus (blue arrows). (B) SIM imaging of untagged P granules (green) immunostained with anti-PGL-1 also reveals P granule pockets (red arrows) indicating fluorescent tags are not influencing P granule morphology. To avoid additional fluorescent tags, *Mutator* foci (magenta) were visualized via SNAP::HA::MUT-16 immunostained with anti-HA. Untagged P granule pockets also associate with *Mutator* foci (blue arrows).

**Figure S3.**
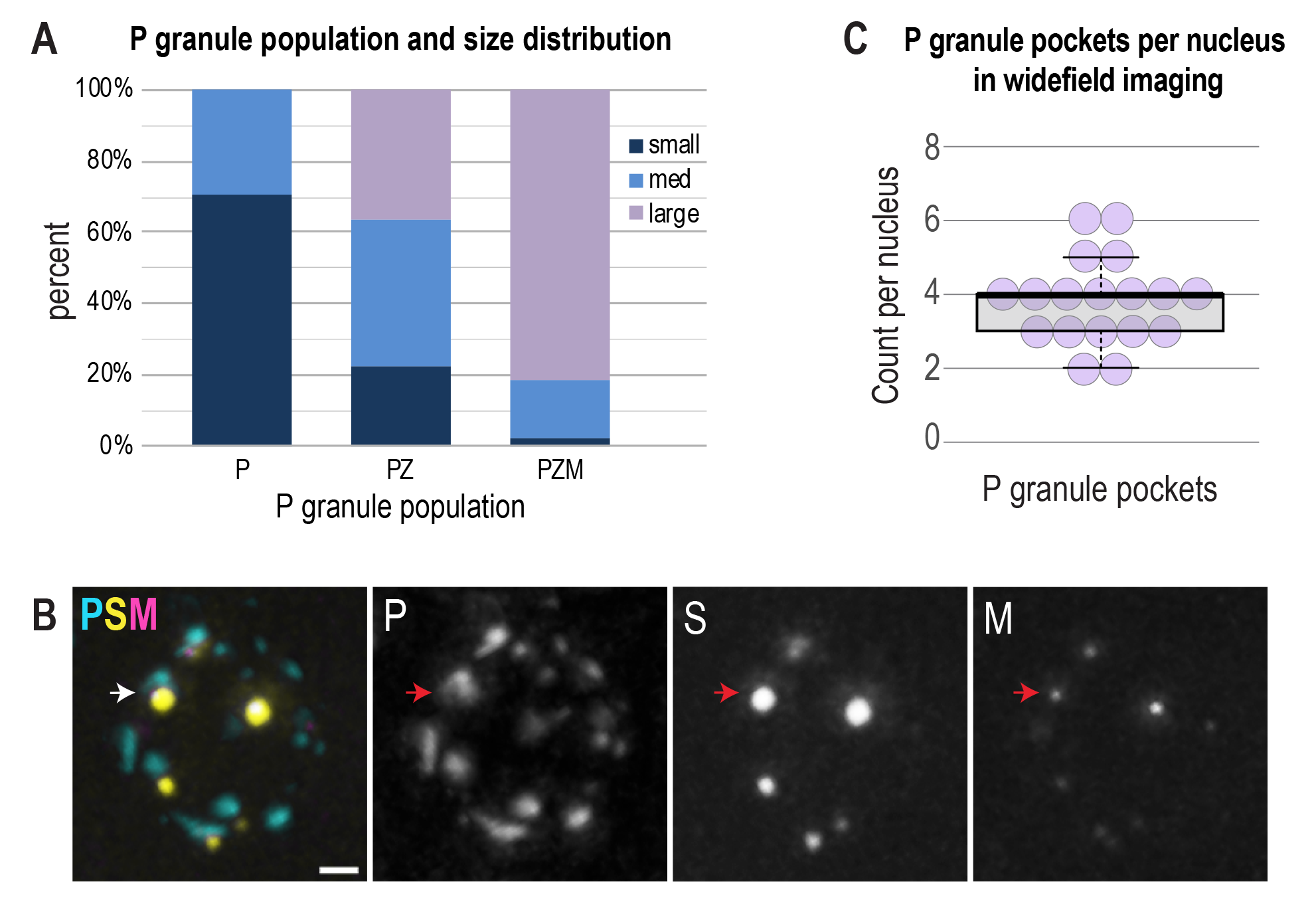
Further analysis of nuage compartment populations. (A) P granules (n = 173) were first qualitatively assessed as small, medium, or large in size and subsequently sorted according to associated nuage compartments (P, PZ, or PZM). 71% of P granules associated with no other compartments were small. P granules associated with Z granules were more evenly distributed in size. 81% of P granules associated with all other granules were large. (B) Representative widefield image of a fixed *simr-1::gfp; rfp::znfx-1; pgl-1::bfp* late pachytene nucleus showing a P granule associated with both SIMR foci and *Mutator* foci (arrow). Scale bar, 1 µm. (C) Box plot showing quantification of P granule pockets per nucleus (n = 18) in the late pachytene where each dot corresponds to one nucleus.

## Notes

### Competing Interest Statement

The authors have declared no competing interest.

